# HIV Coreceptors Related Genes (CCR5 and CXCR4) Promoter Methylation: Original Research

**DOI:** 10.1101/2024.07.01.601546

**Authors:** Anna Esman, Svetlana Salamaikina, Alina Kirichenko, Michael Vinokurov, Darya Fomina, Kirill Sikamov, Arina Syrkina, Anastasiya Pokrovskaya, Vasily Akimkin

## Abstract

The persistence of human immunodeficiency virus (HIV) within viral reservoirs poses significant challenges to eradication efforts. Epigenetic alterations, including DNA methylation, are potential factors influencing HIV latency and persistence. This study details the development and application of techniques to assess CpG methylation in the promoter regions of the *CCR5* and *CXCR4* genes, key HIV-1 coreceptors. Using both Sanger sequencing and pyrosequencing methods, we examined 51 biological samples from 17 HIV-1-infected individuals at three time points: baseline (Week 0) and post-antiretroviral therapy (ART) at Weeks 24 and 48. Our results revealed that *CXCR4* promoter CpG sites were largely unmethylated, while *CCR5* promoter CpGs exhibited significant variability in methylation levels. Specifically, *CCR5* CpG 1 showed a significant decrease in methylation from Week 0 to Week 48, while *CXCR4* CpG 3 displayed a significant decrease between Week 0 and Week 24. These differences were statistically significant when compared with non-HIV-infected controls. These findings demonstrate distinct methylation patterns between *CCR5* and *CXCR4* promoters in HIV-1 positive individuals over time, suggesting that epigenetic modifications may play a role in regulating HIV-1 persistence. Our techniques provide a reliable framework for assessing gene promoter methylation and could be applied to further research in HIV epigenetics.

## 1. Introduction

Lifelong persistence of human immunodeficiency virus (HIV) in infected organisms is a global public health problem. Antiretroviral therapy (ART) significantly reduces mortality and risk of transmission in HIV-individuals through suppression of viral replication [1] [2] [3] [4].

HIV RNA counts reduced in ART recipients with sustained virologic response to undetectable values (<50 copies/mL plasma). However, the provirus activates within a few weeks after the interruption of ART by an increase in viral load [5] [6].

HIV-1 enters into CD4+ T-cells by chemokine coreceptors and persists as a provirus. Recent data indicates that resting CD4+ T-cells are a basic reservoir of the virus and a primary barrier to HIV-1 elimination, despite ART [7]. Current therapy does not cure HIV-infection, and the virus continues to persist in reservoirs.

The chemokine receptors CXCR4 and CCR5 as major coreceptors for HIV-1 entry into CD4+ T-cells. The *CCR5* gene encodes a CCR5 coreceptor for HIV entry into the cell [8] and also influences disease pathogenesis [9]. HIV-1 mainly uses CXCR4 to enter the cell as coreceptor but 5% of virus strains use it as independent receptor [8], [10], [11]. During the early stages of clinical HIV-1 infection predominantly R5-tropic viruses are transmitted. On the contrary, X4-tropic viruses are commonly detected in humans who have received multiple ART regimens and are in the later stages of infection with rapid disease progression [12] [13] [14] [15], [16].

The resting CD4+ T-cells can be roughly divided into two types: naïve ones and HLA-activated (memory) cells. Naïve cells, in contrast to activated cells, predominantly express CXCR4 [17] , while memory cells feature high expression of the *CCR5* gene, which could contribute to HIV replication [18].

Reduced CXCR4 expression affects the HIV-1 life cycle, which is a hallmark of naïve cells that have not passed the activation stage. High expression of the *CCR5* gene affects disease progression and reduces the rate of immune recovery during the course of ART [19]. Reduced expression of the *CCR5* gene correlates with decreased HIV-1 concentration in the cell [20].

The *CCR5* expression level on the surface of activated CD4+ T-cells [17], [18], [21] is increased in HIV-positive individuals [16]. High CCR5 levels could activate resting T-cells, causing apoptosis and their destruction. Therefore, therapy that blocks *CCR5* expression could be associated with slower depletion of the CD4+ T-cells pool without an antiviral effect [22].

Deletion of 32 base pairs of the *CCR5* gene results in a truncated form of the protein that does not fulfill its function. Such deletion contributes to blocking the entry of R5 tropic viruses, forming resistance against nearly 50% of HIV-1 variants [8], [23].

The mechanism underlying the functional effect of methylation in promoter regions could be the disruption of the functioning of regulatory elements at the genomic level caused by changes in the structure of loci and the interaction of transcription factors with DNA [24]. The correlation between decreased expression level and methylation is observed more frequently in the proximity of the transcription initiation site. The main related forms of epigenetic inheritance are genomic imprinting, DNA methylation (CpG loci) and histone modification [25]. Gene methylation regulates its function, leading to a decrease in coreceptor’s gene expression on the cell surface, and this could prevent HIV-1 entry into the cell. Epigenetic modifications, especially methylation, are not linked to DNA sequence changes and play an important role in normal cell activities.

Cytosine methylation as 5mC in CpG loci in the promoter region of genes is associated with transcriptional silencing and plays a central role in epigenetics [26] [27] [28] [29] [30].

Increased DNA methylation in cis-regulatory modules reduces *CCR5* gene activity. There is a significant correlation between the methylation level of the gene and the *CCR5* expression on the surface of T-cells [22].

The increased expression level is associated with demethylation of the cis-modules of *CCR5* gene. Naïve cells are activated by HLA-receptors in response to infection. It has been shown that increased CCR5 expression level can directly support T-cell activation. Hypermethylation is associated with low viral load levels and increased CD4+ T-cell counts; however, cis-modules in HIV-positive individuals remain demethylated compared to normalized CD4+ T-cell counts [22].

*CCR5* gene cis-modules are demethylated and gene expression level rises, which increases the amount of protein on a cell membrane. HIV-1 enters uninfected CD4+ T-cells through CCR5 coreceptor [22].

Suppression of the virus replication by ART has been associated with increased methylation level of *CCR5* [22].

Despite the lack of studies reporting the effect of *CXCR4* gene methylation on HIV-1 replication, there is data that show a strong direct correlation between expression level and the potential HIV-1 entry into the cell [31]. Activation of CD4+ T-cells partly reduces the number of CXCR4 receptors that limit the spread of infection [10]. Reduced levels of the *CXCR4* gene expression correlate with disease progression [16].

In this study we develop techniques and assess the impact of the methylation status in promoter regions of *CCR5* and *CXCR4* genes during ART on HIV-individuals compared to healthy non-HIV infected control.

## 2. Materials and Methods

### 2.1. Study Population

Depersonalized samples from 17 antiretroviral-naïve individuals with HIV-1 infection were collected during 2021–2023 and 27 non HIV-infected individuals during 2024 in Infectious disease clinics of Central Research Institute of Epidemiology (CRIE) used in this study. HIV was diagnosed in accordance with country-specific based on the WHO guidelines: made on the basis of two positive ELISA tests confirmed by immunoblot [32], [33].

Inclusion criteria: age over 18 years, viral load <500,000 copies/mL, CD4+ T-cell count >200 cells/mm^3^, less than 3 visits, written informed consent.

The first line ART regimen prescribed was lamivudine/dolutegravir (3TC/DTG). Whole blood samples from HIV-1 positive individuals were obtained at baseline (Week 0) and after ART started (Weeks 24 and 48). In summary, the study included 51 samples from 17 HIV-1 positive individuals at three visits.

### 2.2. Biological Samples

CD4 T-cell counts were obtained via the flow cytometer FACSCalibur (Becton Dickinson, Franklin Lakes, NJ, USA).

Whole human blood samples were examined for the presence of HIV DNA using the AmpliSens^®^ “DNA-HIV-FL” reagent kit (Amplisens, CRIE, Moscow, Russia). The reagent kit includes a plasmid calibrator with a known concentration of human *β-globin* gene and HIV *pol* gene.

Samples purified with “Hemolytic” reagents (Amplisens, CRIE, Moscow, Russia) are used for selective lysis of blood erythrocytes during pre-processing of clinical material of whole peripheral blood for prioring to DNA extraction.

DNA extraction was carried out by using the “RIBO-prep” kit (Amplisens, CRIE, Moscow, Russia). The collected samples were subjected to lysis using a solution, which resulted in the breakdown of cellular membranes and other biopolymer complexes, releasing nucleic acids and cellular components. After the addition of a precipitation solution and centrifugation, the dissolved DNA precipitated, while other components of the lysed clinical material remained in solution and were removed through subsequent washing. In the final step of the extraction, the precipitate was dissolved in an elution buffer, resulting in purified DNA in solution. This procedure yielded a purified DNA preparation that was free from amplification reaction inhibitors, ensuring high analytical sensitivity for PCR analysis.

The concentration of the starting DNA was measured by real-time PCR for the *β-globin* gene using plasmid calibrators with concentrations of 10,000 and 100 copies per mL. The obtained values were then converted from “copies/μl’’ to “ng” using the SciencePrimer tool-Copy number calculator for real-time PCR [34] (where 1 ng DNA is approximately 300 copies). The average concentration of starting DNA before the bisulfite conversion reaction was — 9000 copies per μl or 30 ng.

Bisulfite conversion was performed on samples using the “EpiTect Bisulfite KIT” (QIAGEN, Hilden, Germany). DNA samples were kept under —20°C after bisulfite conversion and eluated 10x in the TE-buffer (AmpliSens, CRIE, Moscow, Russia) for PCR analysis.

It is necessary to note that incomplete bisulfite conversion of DNA results in the interpretation of unconverted cytosines as methylated in assays [35]. Large amounts of DNA (> 10 μg/ml) could decrease the conversion rate by depleting the available bisulfite in the reaction mix and by increasing the pH due to the formation of NH4+ as a by-product of the reaction.

### 2.3. CpG loci selection

The database of the University of California, Santa Cruz (UCSC) was used to select the *CCR5* and *CXCR4* promoter regions and related predicted transcription factors (JASPAR CORE) [36]. The whole *CXCR4* promoter region was chosen for this study. According to the database *CCR5* gene contains 2 promoter regions: promoter region 1 follows the first exon and produces a truncated product that is primarily expressed in naïve T-cells and in memory T-cells [37] and promoter region 2 contributes to transcription of a product with exon 1 which is typical for T-cells. We studied promoter region 2, since the results of Gornalusse et al. research showed that hypermethylation of this DNA region correlates significantly with *CCR5* expression level [22].

Target loci were chosen by comparing promoter sequences from the EPD (Eukaryotic Promoter Database) [38], which contains experimentally confirmed regions at the beginning of transcription, with sequences from CpG islands, i.e., regions where CpG is present more frequently than in other regions.

Coordinate data and CpG identifiers were obtained from the Illumina GRCh37.p13 genome assembly methylation array (GEO Access Viewer) [39] and NCBI database [40].

A typical procedure for methylation research would include a gene expression study, the identification of genes which are downregulated, and NCBI search (National Center for Biotechnology Information) [40] to locate the promoter sequence and transcription start site of the gene. The selected genomic sequence is then used to aid in the selection of primers.

The detailed description of chosen loci and predicted transcription factors presented in **Table 1** and **Table 2**. CpGs presented in our study indicated by blue colour.

**Table 1.**
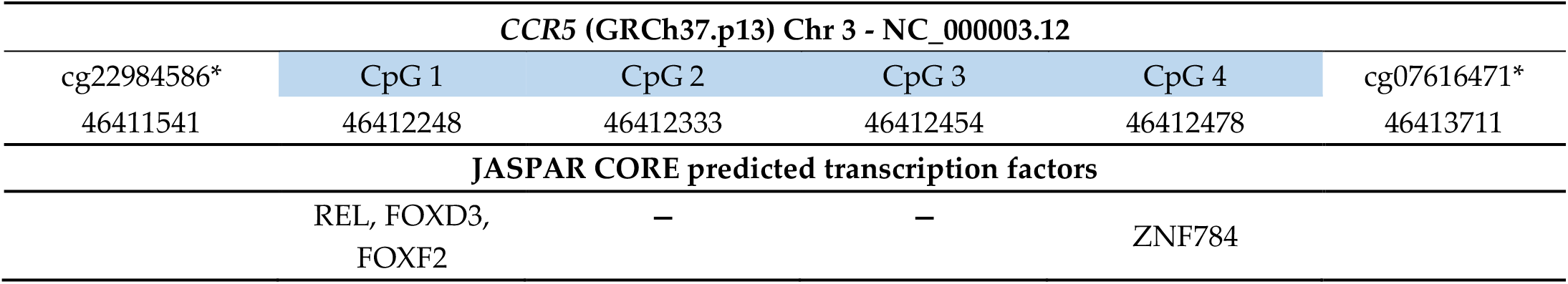
CpG coordinates and predicted transcription factors located in the *CCR5* gene.

**Table 2.**
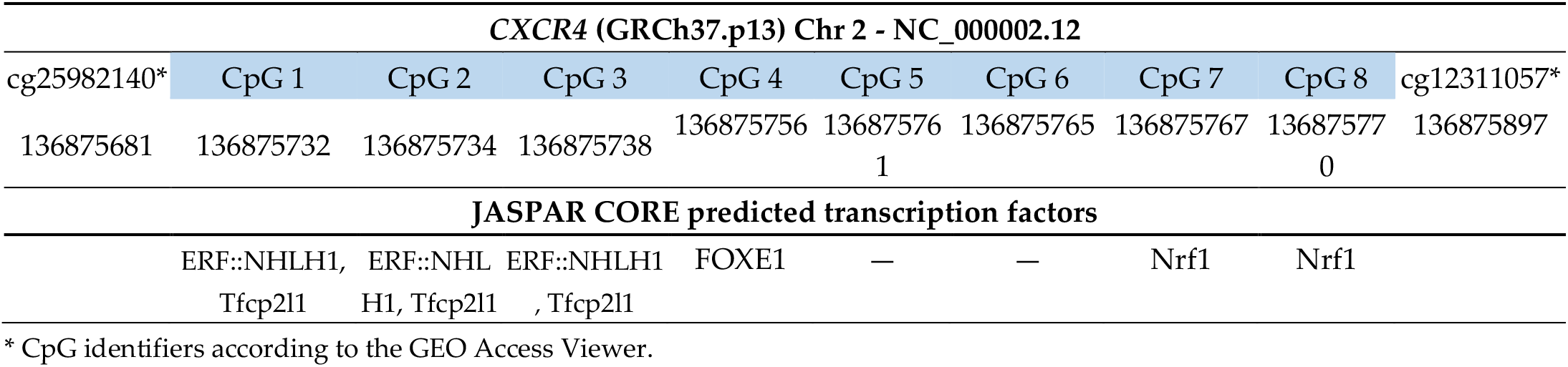
CpG coordinates and predicted transcription factors located in the *CXCR4* gene.

### 2.4. Primers Design

Selected CpG loci were detected by pyrosequencing and Sanger sequencing methods. **Figure 1** shows the common scheme of the oligonucleotide positions.

**Figure 1.**
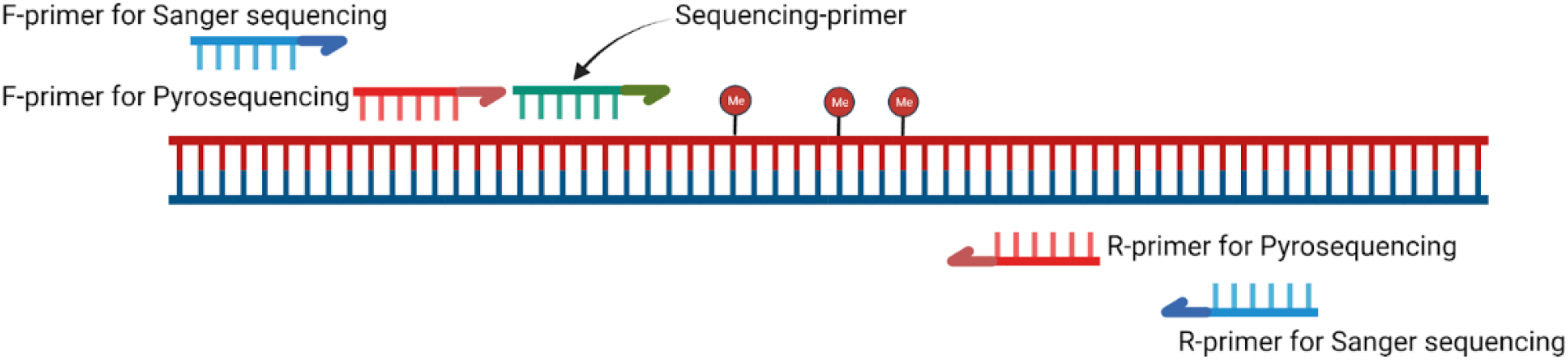
Common scheme of techniques for determining the methylation level of CpG loci (created with BioRender [42]).

*NB!* CpGs at annealing sites should be avoided in oligonucleotide design. The length of the amplicon for pyrosequencing is in the range of 100–250 bp [41].

Oligonucleotides were designed according to standard conditions. The Tm mean calculated in the OligoAnalyzer™ Tool [43] was 60°C. The CG content is rather low because of the features of the converted DNA product and averages 33%. **Table 3** summarizes the sequences of the oligonucleotides used in this study.

**Table 3.**
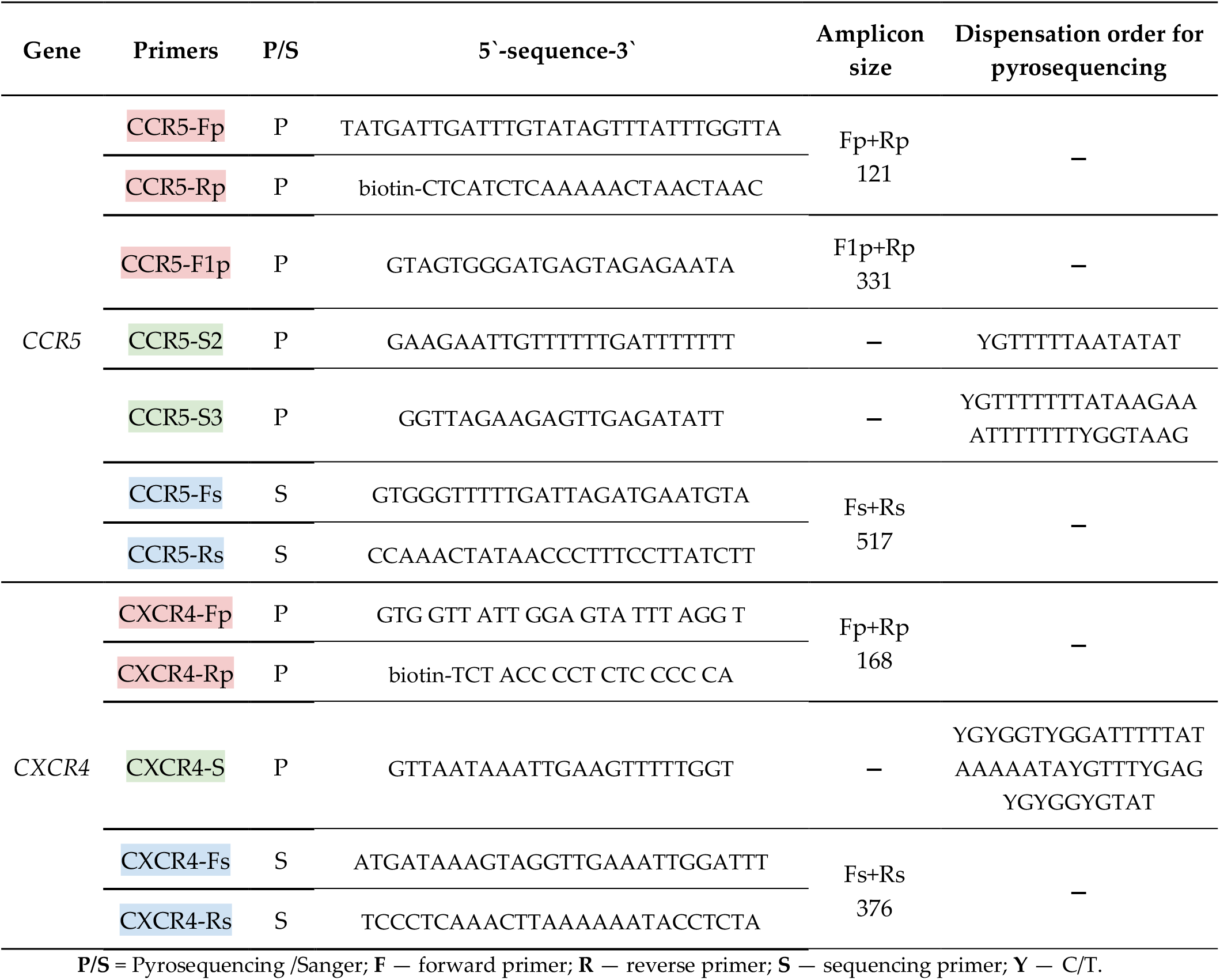
Oligonucleotides used in the study.

**Table 3.**
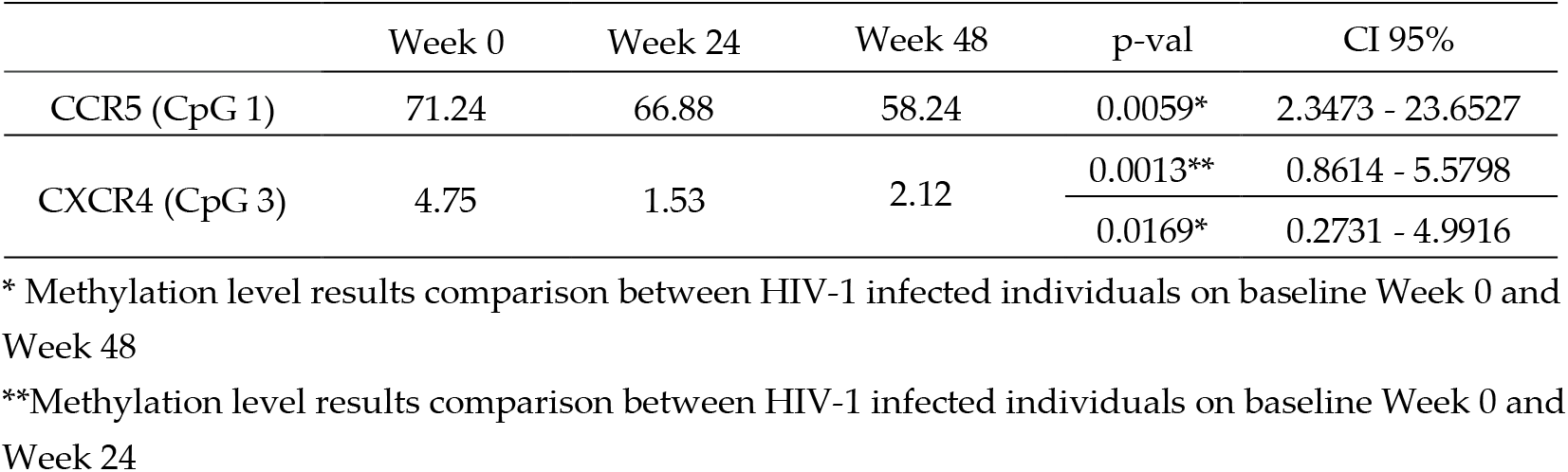
Methylation level differences in HIV-1 positive individuals observed baselines.

|  | Week 0 | Week 24 | Week 48 | p-val | CI 95% |
| --- | --- | --- | --- | --- | --- |
| *CCR5* (CpG 1) | 71.24 | 66.88 | 58.24 | 0.0059* | 2.3473 - 23.6527 |
| *CXCR4* (CpG 3) | 4.75 | 1.53 | 2.12 | 0.0013** | 0.8614 - 5.5798 |
|  |  |  |  | 0.0169* | 0.2731 - 4.9916 |

\* Methylation level results comparison between HIV-1 infected individuals on baseline Week 0 and Week 48
\*\*Methylation level results comparison between HIV-1 infected individuals on baseline Week 0 and Week 24

### 2.5 PCR Conditions

Amplification system “Tercyc” (DNA-technology, Protvino, Russia) was used for PCR sample preparation before the sequencing. The amplification program was as follows: hold 95°C — 15 min, cycling 45 cycles: 95°C — 5 sec, 60°C — 20 sec, 72°C — 5 sec.

PCR reactions were carried out in a final volume of 25 μL, containing 5 μL of PCR-mix-1 (dNTPs and primers); the concentration of primers was experimentally matched — 0.7 μM), 10 μL of PCR-mix-2 (Tris-buffer and MgCl_2_) and 0.5 μL of TaqF polymerase. All used reagents and kits were made in the Central Research Institute of Epidemiology, Federal Service for Surveillance on Consumer Rights Protection and Human Wellbeing Russian Federation (AmpliSens, Moscow, Russia). Finally, 10 μL of bisulfite-treated DNA samples were added to the prepared tubes.

Completely methylated DNA and fully unmethylated DNA, negative control were used as controls for amplification [44], [45] (QIAGEN, Hilden, Germany).

### 2.6. Pyrosequencing Conditions

Sample preparation was carried out with the PyroMark Q24 vacuum station (QIAGEN, Hilden, Germany) and sample preparation kit “Amplisens^®^ Pyroscreen” (Amplisens, CRIE, Moscow, Russia) according to manufacturers’ recommendations.

The reaction was performed with 5 μl of the PCR products. Incubation with sequencing primer was carried out at 80°C for 2 min (at a concentration of 7.5 pmol/μL).

Pyrosequencing was performed using the “PyroMark Gold Q24 Reagents kit” (Qiagen, Hilden, Germany).

Methylation levels of CpGs by pyrosequencing were calculated automatically using PyroMark Q24 software (Qiagen, Hilden, Germany). Quantitative bisulfite Pyrosequencing determines DNA methylation level by analyzing artificial “C/T” SNPs at CpG sites within a specific Pyrosequencing assay [41].

### 2.7. Sanger sequencing Conditions

The amplification system “T100” (BioRad Laboratories, Hercules, USA) was used for PCR sample preparation with the following steps: 96 °C — 1 min, 25 cycles: 96 °C — 10 sec, 50 °C — 5 sec, 60 °C — 4 min. Sanger sequencing was performed using “AmpliSense^®^ HIV-Resist-Seq” reagent kit (Amplisens^®^, CRIE, Moscow, Russia) and Applied Biosystems 3500 genetic analyser (LifeTechnologies, Waltham, USA).

### 2.8. Statistical Methods

Methylation analysis of CpGs by Sanger sequencing was performed according to the protocol of Parrish et al. [46]. Data of methylation level for each CpG loci were preliminarily processed in Microsoft Excel [47]. All statistical tests were run using R 4.2.2 [48], [49]. Descriptive statistical analysis and Two-way ANOVA with Dunnett test (for comparing control non HIV-individuals and HIV-1 positive individuals’ genes methylation level at baseline: Week 0, Week 24 and Week 48) Mann — Whitney U-test (for comparing HIV-1 positive individuals CpGs methylation level at baseline: Week 0, Week 24 and Week 48 with each other) were employed in data evaluation. The results were considered statistically significant at p < 0.01. Data visualization created with Graph application on BioRender.com.

## 3. Results

### 3.1. Sample description

First, we analyzed viral load, viral reservoir size, and CD4+ T-cell counts in samples collected from HIV-infected individuals at baseline (Week 0) and after ART started (Weeks 24 and 48).

Viral load at the first visit for HIV-1 positive individuals varied from 1,371 to 208,538 (copies/mL), median was 38,181 copies/mL. All HIV-1 positive individuals had viral suppression (HIV plasma RNA< 50 copies/mL) at Weeks 24 and 48. There were no treatment-related adverse effects requiring regimen changes. There was no clinical progression of infection in the sense of HIV-associated diseases. All HIV-1 positive individuals reached the virological efficacy of ART by Week 24 of treatment and remained at Week 48.

HIV-1 viral reservoir volume decreased systematically over the course, however there was no statistically significant reduction.

Normally, CD4+ T-cell counts in non HIV-infected individuals range from 500 to 1500 cells per mm^3^. There was a positive trend of CD4+ T-cell counts (**Figure 2**).

**Figure 2.**
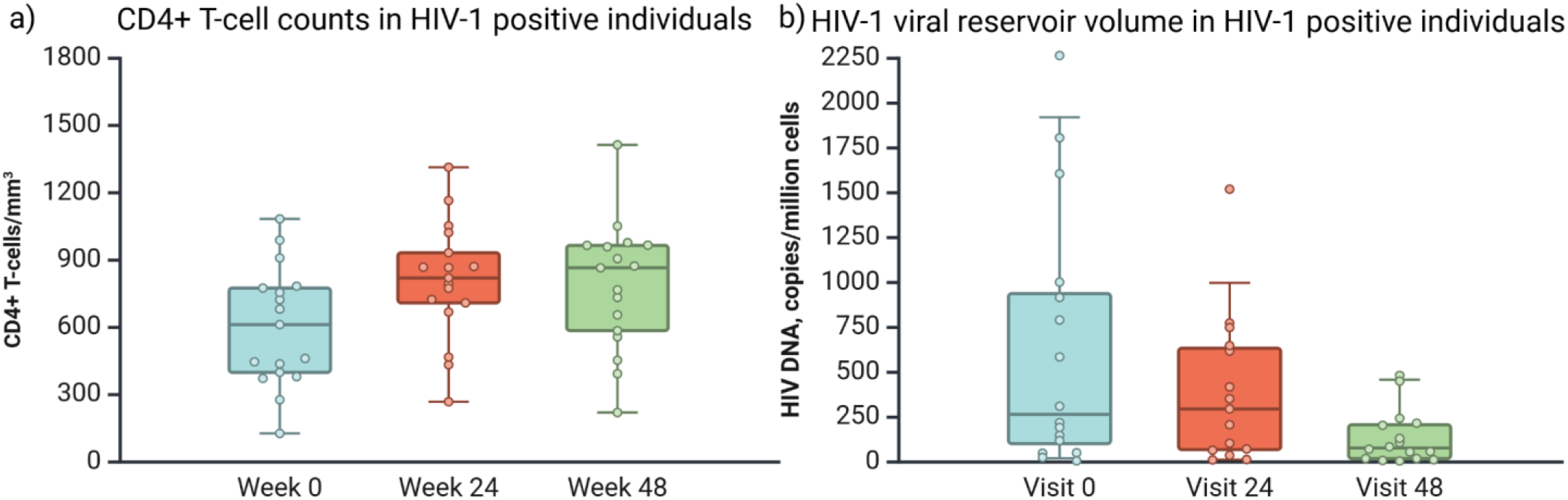
a) CD4+ T-cell counts per mm^3^ and b) HIV-1 viral reservoir volume in samples collected from HIV-1 positive individuals at baseline (Week 0) and after ART (Weeks 24 and 48). Created in BioRender.com [42].

Information about viral load, size of the viral reservoir measured in copies of HIV DNA per million cells, CD4+ T-cell count (cells/mm^3^) for all HIV-1 positive individuals collected in [47].

### 3.2. Development of CpG loci methylation techniques

Pyrosequencing and Sanger sequencing methods were used to assess CpG loci methylation in promoter regions of *CCR5* and *CXCR4* genes. Pyrosequencing is undoubtedly more accurate in measuring the C/(C+T) ratio of peak heights, as this method is the “gold standard” for quantitative allele quantification at single base resolution [50].

**Figure 3.** demonstrates an example of the *CXCR4* gene promoter sequence. Pyrogram on PyroMarkQ24 (Qiagen, Hilden, Germany) in the upper part of the figure shows the same region by Sanger sequencing in the middle and expected sequence in the down part of the figure. Lines link the related CpG loci on the pyrogram and chromatogram.

**Figure 3.**
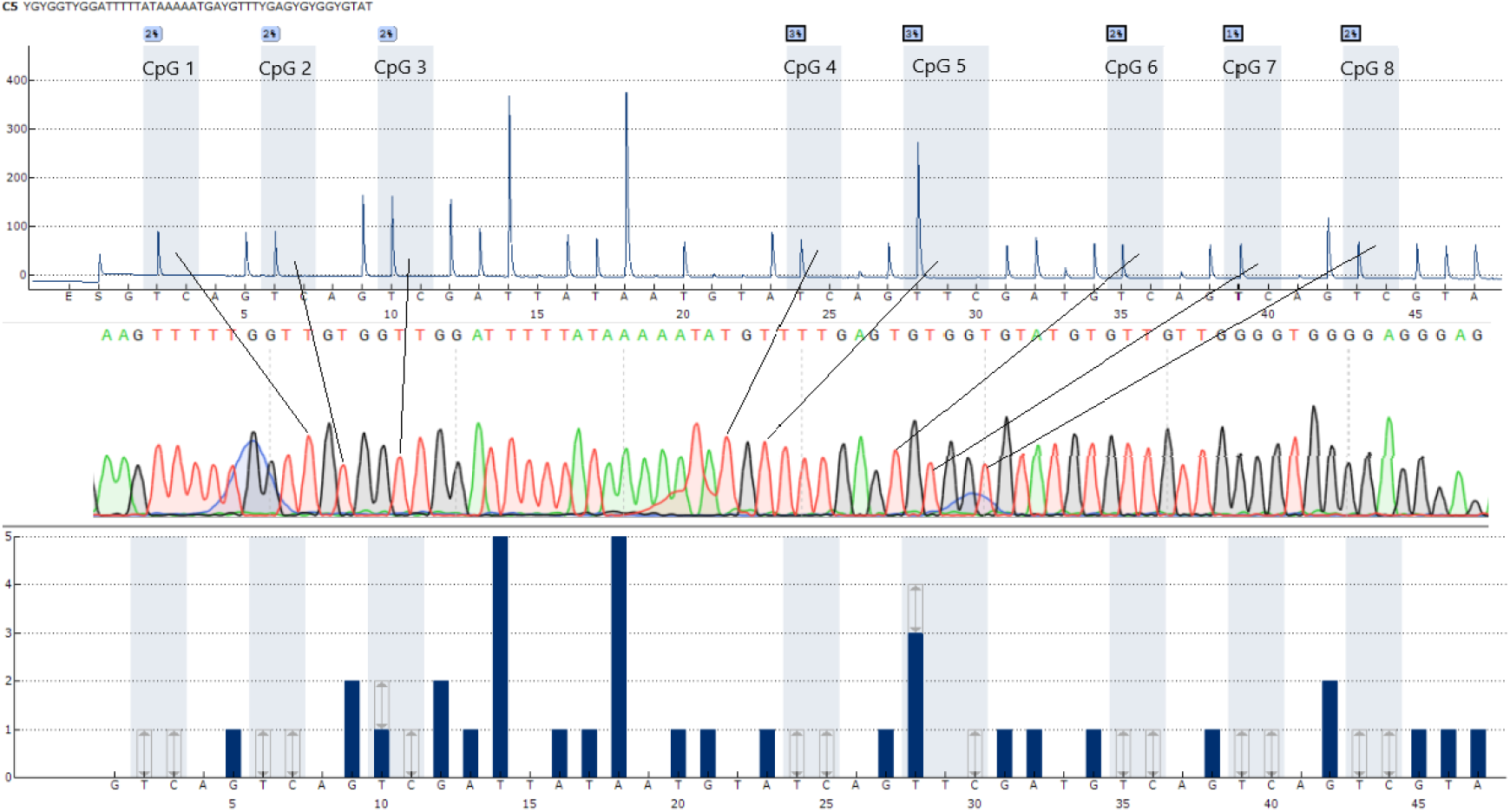
An example of *CXCR4* promoter region sequence.

Considering the CpGs location in the *CCR5* gene second promoter region, several combinations of oligonucleotides were used to read 100–250 bp amplicon length sites by pyrosequencing. These combinations were presented in section “*2.4. Methodology Design/Primers Design*”.

<u>F</u>ully methylated control reaction containing bisulfite-treated DNA (QIAGEN, Hilden, Germany) were on average of 88% and 93.5%, a fully unmethylated control reaction with bisulfite-treated DNA (QIAGEN, Hilden, Germany) were on average of 1% and 4.8% (assays for *CCR5* and *CXCR4* accordingly). The comparison of the results obtained using the pyrosequencing method and Sanger sequencing showed no statistically significant differences.

### 3.3. Application Results of the Developed Techniques

The results obtained using the developed pyrosequencing and Sanger sequencing techniques did not reveal any significant differences. Therefore, for the purposes of this study, we have chosen to use data generated only by the Sanger method.

The mean methylation levels of the *CCR5* and *CXCR4* promoters differences in HIV-1 positive individuals at baseline Week 0, Week 24 and Week 48 and non HIV-infected individuals were statistically significant (p < 0.0001 (**Figure 4**). We also observed that methylation levels of *CXCR4* promoter regions were significantly lower than *CCR5*.

**Figure 4.**
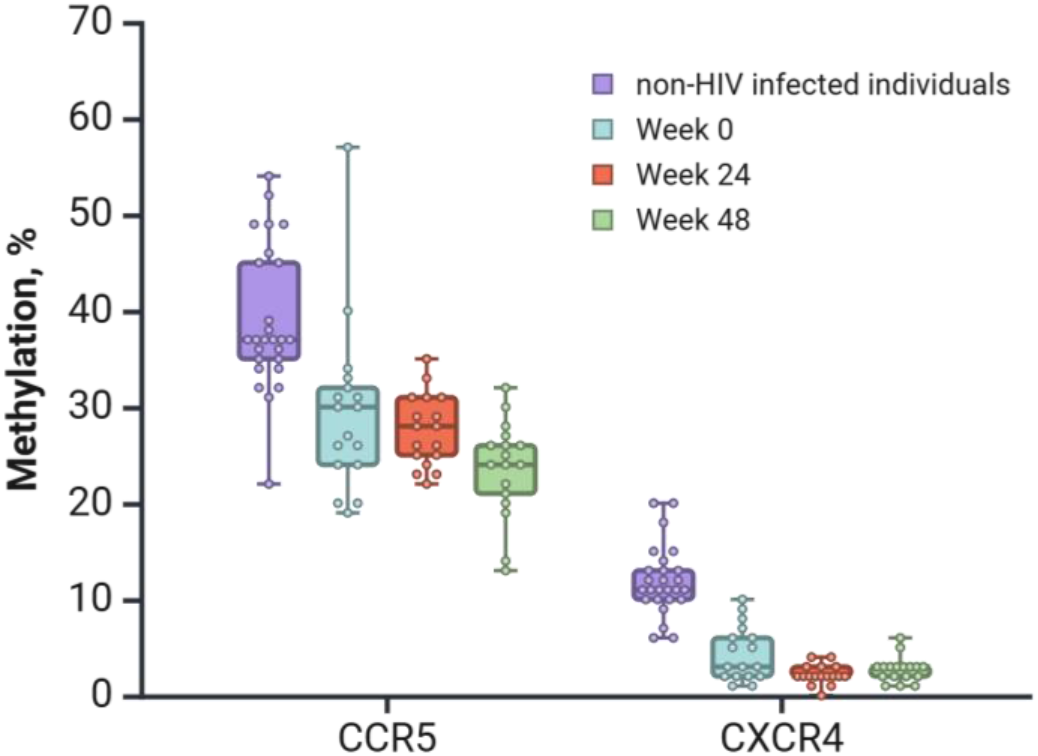
The average methylation level in *CCR5* and *CXCR4* promoter regions in samples collected from HIV-1 positive individuals, assessed at baseline Week 0, Week 24, Week 48 and non-HIV-infected individuals. Created in BioRender.com [42].

Next, we analyzed every CpGs in *CCR5* gene separately in HIV-1 positive individuals at baseline Week 0, Week 24 and Week 48. The CpG 1 methylation level of the CCR5 promoter ranged from 33% to 99%, CpG 2 — from 4% to 83%, CpG 3 — from 0 to 42% and CpG 4 — from 1% to 21%. Analysis of methylation of CpGs in the promoters of the *CCR5* revealed statistically significant differences in methylation levels between HIV-1-infected individuals at different follow-up stages, specifically between the Week 0 and Week 48 groups, at CpG 1 (p = 0.0015) (**Figure 5**).

**Figure 5.**
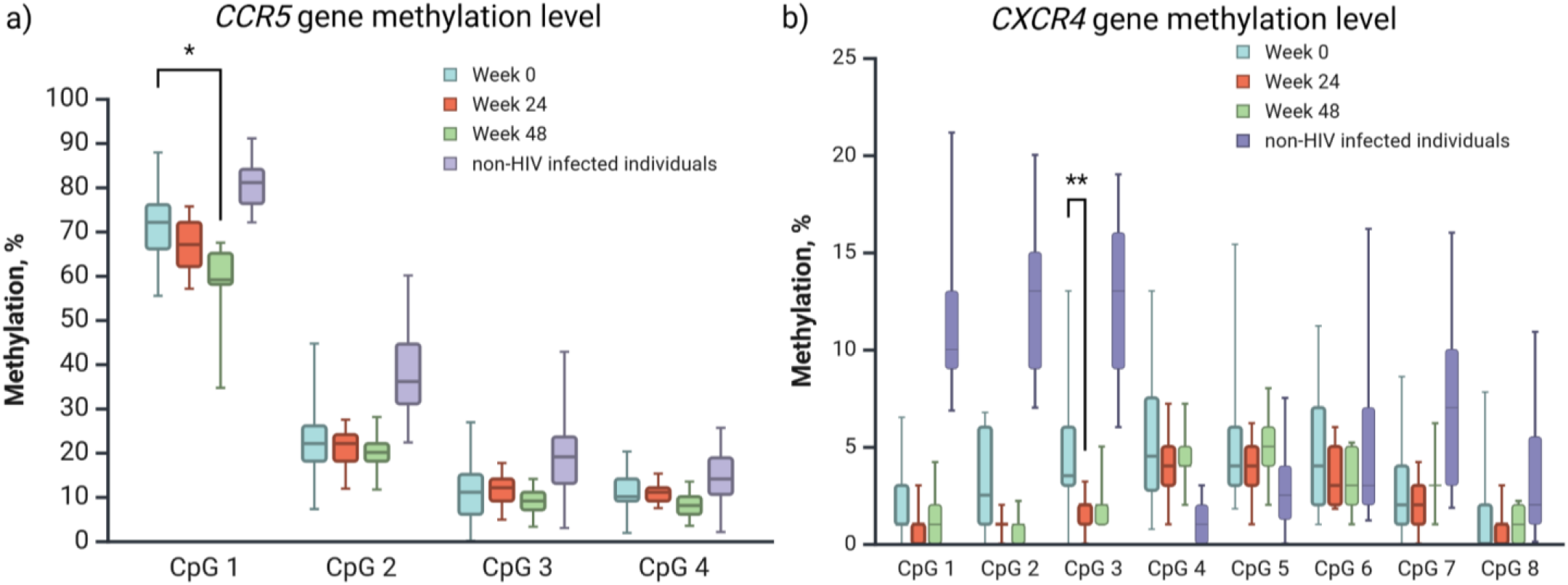
CpGs methylation level in a) *CCR5* and b) *CXCR4* promoter regions in samples collected from HIV-1 positive individuals, assessed at baseline Week 0, Week 24 and Week 48. Created in BioRender.com [42].

Analysis for *CXCR4* promoter region represents CpG 1 methylation level ranged from 0% to 17%, CpG 2 — from 0% to 21%, CpG 3 — from 0% to 21%, CpG 4 — from 0% to 25%, CpG 5 — from 1% to 17%, CpG 6 — from 1% to 16%, CpG 7 — from 0% to 11%, CpG 8 — from 0% to 15%. Statistical differences were observed in CpG3 between baseline Week 0 and Week 24 (p=0.002) (**Figure 5**).

Based on these results, we compared the methylation levels of *CCR5* (CpG 1) and *CXCR4* (CpG 3) CpGs at baselines: Week 0, Week 48 and Week 0, Week 24 accordingly with statistical differences and non-HIV individuals. There were statistical differences observed for the following groups by two-way ANOVA with Bonferroni multiple comparisons test (**Table 3**).

We compared methylation level in promoter regions of *CCR5* and *CXCR4* genes on our group of patients splitted by tropism type. A regular two-way ANOVA was run. There was a significant difference in methylation level of *CXCR4* promoter region between patients with CXCR4 tropism type on baseline Week 24 and patients with CCR5 tropism type on baseline Week 0 (**Figure 6**).

**Figure 6.**
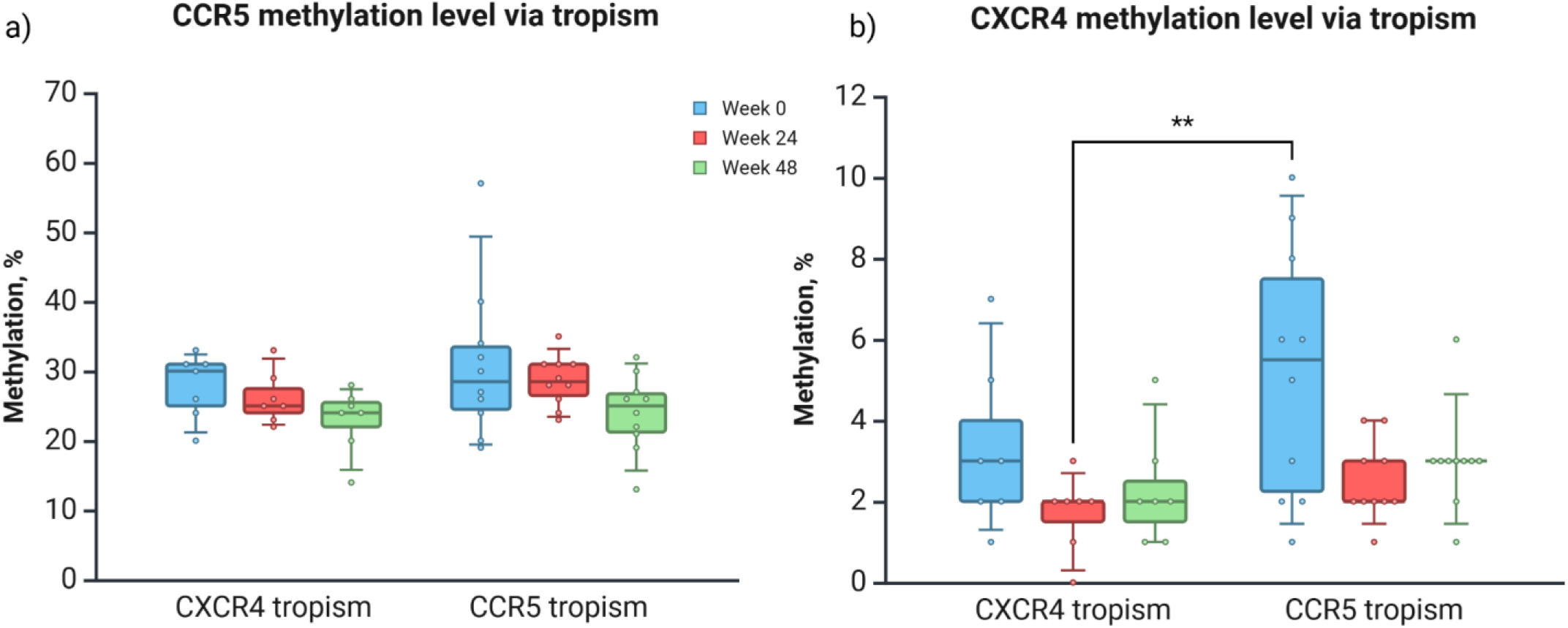
The average methylation level in a) CCR5 and b) CXCR4 promoter regions in samples collected from HIV-1 positive individuals, assessed at baseline Week 0, Week 24, Week 48 and non-HIV-infected individuals splitted by tropism type. Created in BioRender.com [42].

## 4. Discussion

Genome DNA methylation is not uniformly distributed: both promoter and CG-islands typically are hypomethylated, whereas the extent of methylation in gene bodies is higher than that in intergenic regions. While early studies suggested that DNA methylation represses gene expression, a growing body of evidence has indicated that DNA methylation has a dual role, both inhibitory and permissive, depending on the genomic region at which DNA methylation occurs [51].

The association between viral replication and *CXCR4* and *CCR5* expression level suggests a possible direct effect of methylation on these genes on HIV-1 progression within human immune cells [16]. *CCR5* and *CXCR4* receptor genes activity largely regulates the entry of HIV-1 into cells. Gene methylation of the promoter regions decreases the concentration of unintegrated HIV-1 in cells and could contribute to viral eradication [19]. The hypermethylation of *CCR5* and *CXCR4* promoters can reduce both the concentration of HIV-1 RNA inside cells and the risk of infecting rest CD4+ T-cells.

Methylation levels could be analyzed by various methods such as pyrosequencing, which uses bisulfite-treated DNA. However, the Sanger method is more convenient when the required multiple targets are placed more than 60-70 nucleotides apart since a well-read fragment for pyrosequencing is usually limited to about 50 nucleotides. We have demonstrated combinations of primers for pyrosequencing and Sanger sequencing methods in this study. In the case of Sanger sequencing, it will be sufficient to use either one pair of primers or two pairs to suppress the “noise” during result generation.

We also compare these methods on designed primers for detection methylation level of *CCR5* and *CXCR4* promoter regions. Results had no differences between used methods for detection of CpG methylation. While some research groups prefer pyrosequencing as the optimal method, Sanger sequencing also yields satisfactory results [52].

In this study, we have noticed differences in the methylation level between observed CpGs in the *CCR5* promoter region and hypomethylation of the *CXCR4* promoter region. These findings make the study an interesting area for further research.

When DNA methylation affects the promoter region, transcription of the gene is usually suppressed, whereas actively expressed genes tend to have unmethylated promoters. In our data, we found that specific CpGs were more accessible to transcription factors, as identified by JASPAR CORE data [40], where these loci correspond to binding motifs for transcription factors listed in Tables 1 and 2. This suggests that differential methylation at these loci may influence gene expression by modulating the recruitment of these specific transcription factors.

For example, transcription factors predicted to bind in the *CCR5* promoter region, such as REL, a member of the NF-κB/Rel family, plays a critical role in the maintenance of HIV-1 latency, suggesting that methylation at its binding sites could influence the ability of HIV-1 to persist in a latent state [53]. Similarly, FOXD3 acts as a negative regulator of HIV-1 replication in mononuclear phagocytes by suppressing genes involved in viral replication, such as OTK18. The upregulation of FOXD3 during HIV infection could explain the restricted replication of the virus in certain immune cells like microglia [54].

Furthermore, Tfcp2l1 is a member of the Grainyhead-like transcription factor family, which regulates both cell cycle progression and HIV-1 genes expression [55]. Its binding motifs in the promoter regions of *CXCR4* could play a role in controlling the expression of these coreceptors, thus impacting HIV-1 entry into host cells. Additionally, Nrf1 plays a crucial role in regulating *CXCR4* promoter activity, and its involvement could lead to enhanced *CXCR4* expression during immune activation, potentially contributing to increased HIV-1 replication and disease progression [56].

Although we observed some statistically significant differences in methylation levels between HIV-1 positive individuals and non HIV controls. These results could be caused by random variation, considering the small sample size in our study and other factors that may influence methylation levels, such as age, status of other genes etc. Although we observed differences in methylation levels of *CCR5* and *CXCR4* promoter regions, we could not clearly conclude whether these results have a protective effect. However, we can only hypothesize that the activity of *CCR5* gene is reduced in the analyzed samples.

The confirmation of the obtained results, further investigation of the expression levels of these genes is necessary.

The study of gene methylation presents an opportunity to identify potential targets for HIV treatment through the use of genomic editing techniques, which can be combined with ART or potentially replace current treatment regimens [57]. For example, the CRISPR/Cas9 method would allow the delivery of methyltransferases and demethylases to CpG loci to change the methylation status and, consequently, the level of gene expression [58].

Epigenetic modification study of the chemokine receptor group could reveal the connection between HIV tropism and methylation levels of CpGs in promoters, which could further uncover new pathways of HIV elimination. The eradication of HIV-1 in infected people will be impossible without solving the problem of the persistence of the virus in its reservoirs.

## Supplementary Materials

The following supporting information can be downloaded at: https://figshare.com/s/2fce38ebe705a8501770, Table S1: DTG 3TC HIV EPI study.xlsx.

## Author Contributions

Conceptualization, A.E., M.V., S.S. and D.F.; methodology, A.E. and M.V.; validation, S.S., D.F. and A.S.; resources, A.K. and A.P., writing—original draft preparation, A.E., S.S., M.V., D.F., A.K., K.S., writing—review and editing, A.E. and S.S., visualisation, S.S. and A.E., supervision, V.A., project administration, A.E., funding acquisition, A.E. All authors have read and agreed to the published version of the manuscript.

## Funding

This research was funded by Central Research Institute of Epidemiology, grant number 122053000056-2.

## Institutional Review Board Statement

This study was conducted in accordance with the Declaration of Helsinki, and approved by the Ethics Committee of the CRIE, Federal Service for Surveillance on Consumer Rights Protection and Human Wellbeing, Russian Federation (protocol code No 117, 28 September 2021).

## Informed Consent Statement

Informed consent was obtained from all subjects involved in this study.

## Acknowledgments

The authors are grateful to PhD (Med.Sci.) **K. Mironov**, the Head of the Laboratory of Molecular Methods for Genetic Polymorphisms Research, Central Research Institute of Epidemiology (CRIE), PhD (Biol. Sci.) **D. Kireev**, the Head of the Laboratory of Diagnostics and Molecular Epidemiology of HIV Infection (the above Institute) and the senior researcher of the said Laboratory, PhD (Biol. Sci.). **I. Lapovok** for consultations and support with reagents.

## Conflicts of Interest

The authors declare no conflict of interest.

## Abbreviations

The following abbreviations are used in this manuscript:

HIV: Human immunodeficiency virus
ART: Antiretroviral therapy
CRIE: Central Research Institute of Epidemiology
DNA: Deoxyribonucleic Acid
PCR: Polymerase chain reaction
ELISA: Enzyme Linked Immunosorbent Assay
UCSC: University of California, Santa Cruz
EPD: Eukaryotic Promoter Database
NCBI: National Center for Biotechnology Information
ANOVA: Analysis of variance

## Notes

https://figshare.com/s/2fce38ebe705a8501770

### Competing Interest Statement

The authors have declared no competing interest.

### Summary of Updates

We have changed the figures Figure 5. CpGs methylation level in a) CCR5 and b) CXCR4 promoter regions in samples collected from HIV-1 positive individuals, assessed at baseline Week 0, Week 24 and Week 48. Created in BioRender.com [42]. Figure 6. The average methylation level in a) CCR5 and b) CXCR4 promoter regions in samples collected from HIV-1 positive individuals, assessed at baseline Week 0, Week 24, Week 48 and non-HIV-infected individuals splitted by tropism type. Created in BioRender.com [42].

